# ER stress-linked autophagy stabilizes apoptosis effector PERP and triggers its co-localization with SERCA2b at ER-plasma membrane junctions

**DOI:** 10.1101/742882

**Authors:** Samantha J. McDonnell, David G. Spiller, Michael R. H. White, Ian A. Prior, Luminita Paraoan

## Abstract

Specific molecular interactions that underpin the switch between ER stress-triggered autophagy-mediated cellular repair and cellular death by apoptosis are not characterized. This study reports the unexpected interaction elicited by ER stress between the plasma membrane (PM)-localized apoptosis effector PERP and the ER Ca^2+^ pump SERCA2b. We show that the p53 effector PERP, which specifically induces apoptosis when expressed above a threshold level, has a heterogeneous distribution across the PM of un-stressed cells and is actively turned over by the lysosome. PERP is upregulated following sustained starvation-induced autophagy, which precedes the onset of apoptosis indicating that PERP protein levels are controlled by a lysosomal pathway that is sensitive to cellular physiological state. Furthermore, ER stress stabilizes PERP at the PM and induces its increasing co-localization with SERCA2b at ER-PM junctions. The findings highlight a novel crosstalk between pro-survival autophagy and pro-death apoptosis pathways and identify, for the first time, accumulation of an apoptosis effector to ER-PM junctions in response to ER stress.

## Introduction

The p53 apoptosis effector related to PMP-22 (PERP) is a tetraspan plasma membrane (PM) protein that is involved in cell-cell adhesion and the regulation of apoptosis in many cell types.^1–3^ PERP transcription is tightly controlled by both p53 and p63 and is specifically induced during p53-mediated apoptosis.^1,4^ Expression of PERP above a threshold level correlates with the activation and stabilization of its regulator p53 and the cleavage of both caspase 8 and Bid; thus PERP positively influences its own expression and mediates apoptosis engaging both the extrinsic and mitochondrial pathways.^5,6^ PERP is therefore likely to be a key molecule that drives damaged cells towards apoptosis.^7^

Protein synthesis and modification in the endoplasmic reticulum (ER) are highly dependent on a high luminal Ca^2+^ concentration. In addition, spatiotemporal Ca^2+^ signals, generated via release from the ER stores and influx from the extracellular space, regulate many cellular processes including autophagy and cell death.^8^ Sarco/endoplasmic reticulum ATPase (SERCA) pumps regulate Ca^2+^ influx into the ER and are vital for the maintenance of intracellular Ca^2+^ homeostasis. The three SERCA genes (ATP2A1-3) undergo alternative splicing to generate a family of isoforms with tissue-specific expression.^9,10^ SERCA2 is expressed ubiquitously and is recruited to points of contact between the ER and PM involved in the fundamental store operated calcium entry (SOCE) influx mechanism, where it ensures efficient ER Ca^2+^ refilling following depletion.^10,11^

Disruption to ER Ca^2+^, redox and ATP homeostasis results in impaired protein folding and the accumulation of proteins in the ER.^12^ Cell repair is promoted by ER stress signalling pathways which alleviate the damage by inhibiting protein translation, upregulating protein folding and inducing protein degradation.^13^ In addition, increased expression of SERCA enhances ER Ca^2+^ loading in response to both Ca^2+^ depletion from the ER and Ca^2+^-independent ER stress stimuli.^14–16^ Multiple ER stress effectors also rapidly induce autophagy, which degrades protein aggregates and damaged organelles to aid cell survival.^17,18^

In the event of irreparable or prolonged damage, both the ER stress and autophagy pathways interact with those of apoptosis and cell fate is determined by the most dominant response.^19^ In addition, ER Ca^2+^ signalling at regions in close contact with mitochondria has a fundamental role in the regulation of cell fate; high levels of Ca^2+^ transfer results in the loss of mitochondrial integrity, thereby promoting caspase-dependent apoptosis.^20,21^ Toxic levels of intracellular Ca^2+^ are provided by the action of ER and PM pumps/channels.^22–24^ For example, altered SERCA activity mediates apoptosis via ER-mitochondrial Ca^2+^ transfer.^25–28^

This study was initiated by the characterization by mass spectrometry of the protein-protein interactions of PERP required for apoptosis induction. We discovered that PERP is stabilized at the PM due to a reduction in its lysosomal uptake and degradation during ER stress-induced autophagy. Subsequently there is an increased co-localization with the ER Ca^2+^ transporter SERCA2b that correlates with apoptosis induction. The findings thus identified a crosstalk between autophagy and apoptosis pathways with a role in the Ca^2+^-mediated regulation of cell fate at ER-PM junctions.

## Materials and Methods

### Cell culture

Authenticated Mel202 cells were purchased from Public Health England (lot number 13H016) and were cultured in RPMI 1640 with 2mM L-glutamine and 25 mM HEPES (Gibco, Life Technologies, Paisley, UK) supplemented with 10% FCS (Sigma-Aldrich, Dorset, UK), 1 mM sodium pyruvate and 1% non-essential amino acids (Sigma-Aldrich). HCT116 and HCT116 p53−/− cells (obtained from Johns Hopkins University GCRF Core Cell Center, Baltimore, USA; HCT116 p53+/+ (parent of p53 KO), lot 40-16; HCT116 p53−/−, lot 379.2) were grown in Modified McCoy’s 5a medium (Gibco) supplemented with 10% FCS. HeLa cells (original lot purchased from ATCC, catalogue number ATCC CCL-2) stably expressing Venus-PERP from a bacterial artificial chromosome (HeLa BAC Venus-PERP cells, Raheela Awais, personal communication) were cultured in Eagle’s Minimum Essential Medium (ATCC, Middlesex, UK) supplemented with 10% FCS and 0.8 mg/ml geneticin (Sigma-Aldrich). All cells were cultured at 37°C and 5% CO_2_ in humidified incubators and were free of mycoplasma contamination.

### Expression vectors

GFP-PERP, eGFP-SERCA2b and mCherry-SERCA2b were described previously.^6,11^ To generate the Halo-PERP construct, PERP cDNA was amplified by PCR from the GFP-PERP vector using primers designed to add flanking EcoRI and NotI restriction sites to the 5’ and 3’ regions respectively, for subsequent incorporation into the pHTN HaloTag vector (Promega, Southampton, UK). The primer sequences were: forward primer (5’-AATTA**GAATTC**ATGATCCGCTGCGGCCTG-3’) and reverse primer (5’-AATTA**GCGGCCGC**<u>TTA</u>GGCAGATGTGTAGAAGTACCTGGG-3’, with restriction sites shown in bold and stop codon underlined. PCR amplification was performed using the Phusion High-Fidelity PCR Kit (New England Biolabs Inc, Ipswich, UK) and the final reaction contained 250 ng of template DNA, 0.5 μM forward and reverse primers, 200 μM dNTPs, 1X HF buffer and 1 unit of T4 DNA polymerase. Cycling conditions used were 1 cycle of 98°C for 30 seconds, followed by 30 cycles of 98°C for 10 seconds, 63°C for 30 seconds and 72°C for 30 seconds, terminated with one cycle of 72°C for 7 minutes.

Amplified DNA was purified using the QIAquick PCR Purification Kit (Qiagen Ltd, Manchester, UK) and both the PERP sequence and HaloTag vector were digested using EcoRI and NotI (Roche, West Sussex, UK). Ligation was performed using 1 unit of T4 DNA ligase, 1X ligation buffer, 80 ng of backbone DNA and 35 ng of insert DNA. The resulting vector was used to transform DH5α *E. coli* (Invitrogen, Paisley, UK), selected using ampicillin (100 μg/ml) and purified using the Qiagen EndoFree Plasmid Maxi Kit. The sequence of HaloTag-PERP (Halo-PERP) was confirmed by DNA sequencing, performed by DNA Sequencing & Services (School of Life Sciences, University of Dundee, Scotland) on an Applied Biosystems model 3730 automated capillary DNA sequencer.

### Cell transfection and treatment

Mel202 cells were transiently transfected with Halo-PERP, mCherry-SERCA2b and control vectors. Cells were seeded at a density of 4×10^5^ cells/well in 6 well plates and the following day were transfected with 1 μg of plasmid DNA and 2 μl of TurboFect *in vitro* transfection reagent (Thermo Scientific, Paisley, UK). For live cell imaging, 3×10^5^ HeLa BAC Venus-PERP cells were seeded in 35 mm glass bottom dishes (Greiner Bio-One, Stonehouse, UK) and transfected with 1 μg of mCherry-SERCA2b using 2 μl of TurboFect. For HaloTag pull-down, 2.5×10^6^ cells were seeded in 100 mm dishes (4 dishes per transfection) and transfected using 5 μg plasmid DNA and 10 μl of TurboFect. Cells were treated with 1 μg/ml BFA, 20 μM MG132, 100 μM chloroquine, or 1,10 and 100 nM bafilomycin A1 (all from Sigma-Aldrich).

### HaloTag protein pull-down

PERP protein-protein interactions were isolated using the HaloTag Mammalian Pull-Down System (Promega) and standard protocol. Briefly, 1×10^7^ Mel202 cells were transfected with either HaloTag or Halo-PERP plasmids and cell lysates were collected 24 h post-transfection. A total of 2 mg of each protein lysate was incubated with pre-washed HaloLink resin at 4°C overnight. The resin was washed 5 times using TBS with 0.05% IGEPAL CA-360 (Sigma-Aldrich) and PERP plus any interacting proteins were cleaved from the resin using 30 units of ProTEV enzyme (Promega) at room temperature for one hour (elution 1). Following removal of the protein-containing supernatant, the resin was heated to 95°C for 5 minutes in SDS sample buffer (elution 2). Proteins in both elutions were identified by mass spectrometry and independently confirmed by Western blot.

### Immunoprecipitation

SERCA2 was immunoprecipitated using the Dynabeads Protein A Immunoprecipitation Kit (Invitrogen) and monoclonal SERCA2 antibody (1:100, ab2861 Abcam, Cambridge, UK). Cells were scraped in dPBS, pelleted and frozen to −80°C. The resulting pellet was lysed in Mammalian Lysis Buffer with Protease Inhibitor Cocktail (both from Promega) and 1.5 mg of protein was incubated with the bead-antibody conjugate for 30 minutes at room temperature. Antibody-protein complexes were washed 4 times in PBS and proteins were removed in SDS sample buffer for Western blot analysis.

### In-gel digestion and mass spectrometry

For mass spectrometry analysis, proteins were separated on NuPAGE 4-12% Bis-Tris gradient gels (Invitrogen) and each lane was cut into three slices. Samples were reduced in 10 mM dithiothreitol at 56°C, followed by alkylation using 50 mM iodoacetamide at room temperature for 30 minutes. Proteins were digested using 0.08 μg Trypsin Gold (Promega) per gel slice at 37°C for 16 h and peptides were extracted from the band pieces using acetonitrile, dried in a SpeedVac and re-suspended in 1% formic acid.

A total of 5 μl of peptides were separated on a nanoACQUITY UPLC system (Waters, Herts, UK), followed by a LTQ. Orbitrap XL mass spectrometer (Thermo Scientific) with a Proxeon nanoelectrospray source. Peptides were passed through a 5 cm × 180 μm BEH-C18 symmetry trapping column, followed by a 25 cm × 75 μm BEH-C18 column (both from Waters) at a flow rate of 400 nl/min for 39 minutes. The mass spectrometer acquired full scan MS spectra (m/z 300-2000, resolution 30 000) and the five most abundant ions were further fragmented for MS/MS analysis in the LTQ. A blank of 1% formic acid was ran between each sample. Raw mass spectrometry files were analysed using Maxquant proteomics software (version 1.5.3.30) and proteins were identified using the whole HumanIPI database (June 2016).

### Super resolution live cell microscopy

Super resolution imaging was performed using a Zeiss LSM 880 Axio Observer microscope with Airyscan detection system in SR mode using a Plan-Apochromat 63x/1.4 Oil DIC M27 objective (Carl Zeiss, Jena, Germany). Excitation was achieved using the 488nm line from an argon laser and a 561nm diode laser or 594nm HeNe laser and emitted light was collected through appropriate filters to eliminate any spill-over. Raw images were immediately Airyscan processed using the default settings and analysed using ZEN 2.1 software.

### Fluorescent microscopy

Halo-PERP was labelled with the TMRDirect HaloTag ligand and 24 h post-transfection cells were imaged using a Zeiss Axio Observer Z1 live cell microscope with Apotome.2 system and a 40x/1.3 Oil DIC objective (Carl Zeiss).

### Western blotting and antibodies

Cells were harvested following transfection or treatment at specific time points in lysis buffer (0.128 M β-mercaptoethanol, 40 mM tris, 10% glycerol, 1% SDS and 0.01% bromophenol blue) containing PhosSTOP phosphatase and Complete protease inhibitor cocktails (both from Roche). Proteins were separated by SDS-PAGE on a 10% polyacrylamide gel, transferred to nitrocellulose membrane and probed with the primary antibody at 4°C overnight. Antibodies for PERP (ab5986), SERCA2b isoform specific (ab137020) and GAPDH (ab8245) were purchased from Abcam and antibodies for LC3B (#3868) and p62 (#8025) were from Cell Signalling Technology (Leiden, The Netherlands). Anti-HaloTag (G9211) was from Promega and anti-p53 (P6874) was from Sigma-Aldrich. Immunocomplexes were incubated with the appropriate horseradish peroxidase-conjugated secondary antibody and detected using SuperSignal West Pico Chemiluminescent Substrate (Thermo Scientific) or Radiance Plus Chemiluminescent Substrate (Cambridge Bioscience, Cambridge, UK) and a Bio-Rad Chemidoc Imaging System. Membranes were incubated in stripping solution at 55°C for 25 minutes and sequentially re-probed.

### RNA extraction, reverse transcription and RT-PCR

RNA was extracted using an RNeasy Mini Kit (Qiagen) and converted to cDNA using the First Strand cDNA Synthesis Kit (Thermo Scientific). Real-time quantitative PCR was performed for both PERP and GAPDH using the primers and protocol described previously.^29^ Primers were designed to specifically amplify the full length SERCA2 transcript (SERCA2b) at an annealing temperature of 60°C (forward primer 5’-TCGAACCCTTGCCACTCATC-3’ and reverse primer 5’-GCACAAACGGCCAGGAAATC-3’, synthesised by Eurogentec, Southampton, UK).

### Flow cytometry apoptosis detection

Floating and attached cells were collected and incubated with Alexa Fluor 647 annexin V Ready Flow Conjugate (Thermo Scientific) for 15 minutes and 10 000 cells were analysed using a BD Accuri C6 flow cytometer.

### Statistical analysis

The data presented represents the mean ± SEM of at least three independent biological experiments. Data from each biological replicate was assessed for variance within the respective group and only data with similar variance between groups was included in the statistical analysis. One-way ANOVA with Dunnett’s post hoc test, two-way ANOVA with Bonferroni post hoc test, or Student’s t-test were used where appropriate to compare data to the control group. *p<0.05, **p<0.01, ***p<0.001, ****p<0.0001.

## Results

### Identification of PERP – SERCA2b interaction and co-localization

To identify novel effectors involved in PERP-mediated apoptosis we investigated the protein-protein interactions of PERP using the HaloTag Mammalian Pull-Down System. A construct containing PERP cDNA fused to an N-terminal HaloTag was generated and expressed in the human uveal melanoma cell line Mel202 where the fusion protein localized to the PM and secretory pathway (Fig. S1 A), as previously characterised.^6^ Expression of Halo-PERP for 48 h significantly increased the protein levels of endogenous PERP and p53 and induced phosphatidylserine externalisation (Fig. S1 B and C), in line with previous results showing that Halo-PERP functions in both p53 stabilization and apoptosis induction.^5^ This data validated the HaloTag system as a suitable platform for characterisation of the PERP protein interactome.

Halo-PERP protein pull-down and subsequent mass spectrometry analysis identified 21 proteins with a fold change of 1.5 or higher in at least two independent experiments and 6 proteins were identified exclusively in Halo-PERP pull-downs in all three experiments. Interestingly, 3 ER membrane proteins were repeatedly identified in Halo-PERP pull-downs and were absent from all controls. Of these, we chose to focus on SERCA2 (Fig. 1 A). The SERCA2 gene has three isoforms, SERCA2a-c, which differ in their C-terminal region and tissue-specific expression. SERCA2b is the house-keeping isoform which has the highest affinity for Ca^2+^ and is expressed in all tissues.^10^ As the peptide identified by mass spectrometry could not distinguish between the three isoforms, the specific interaction between PERP and SERCA2b was confirmed after HaloTag pull-down by Western blot using the isoform specific antibody (Fig. 1 B).

**Figure 1.**
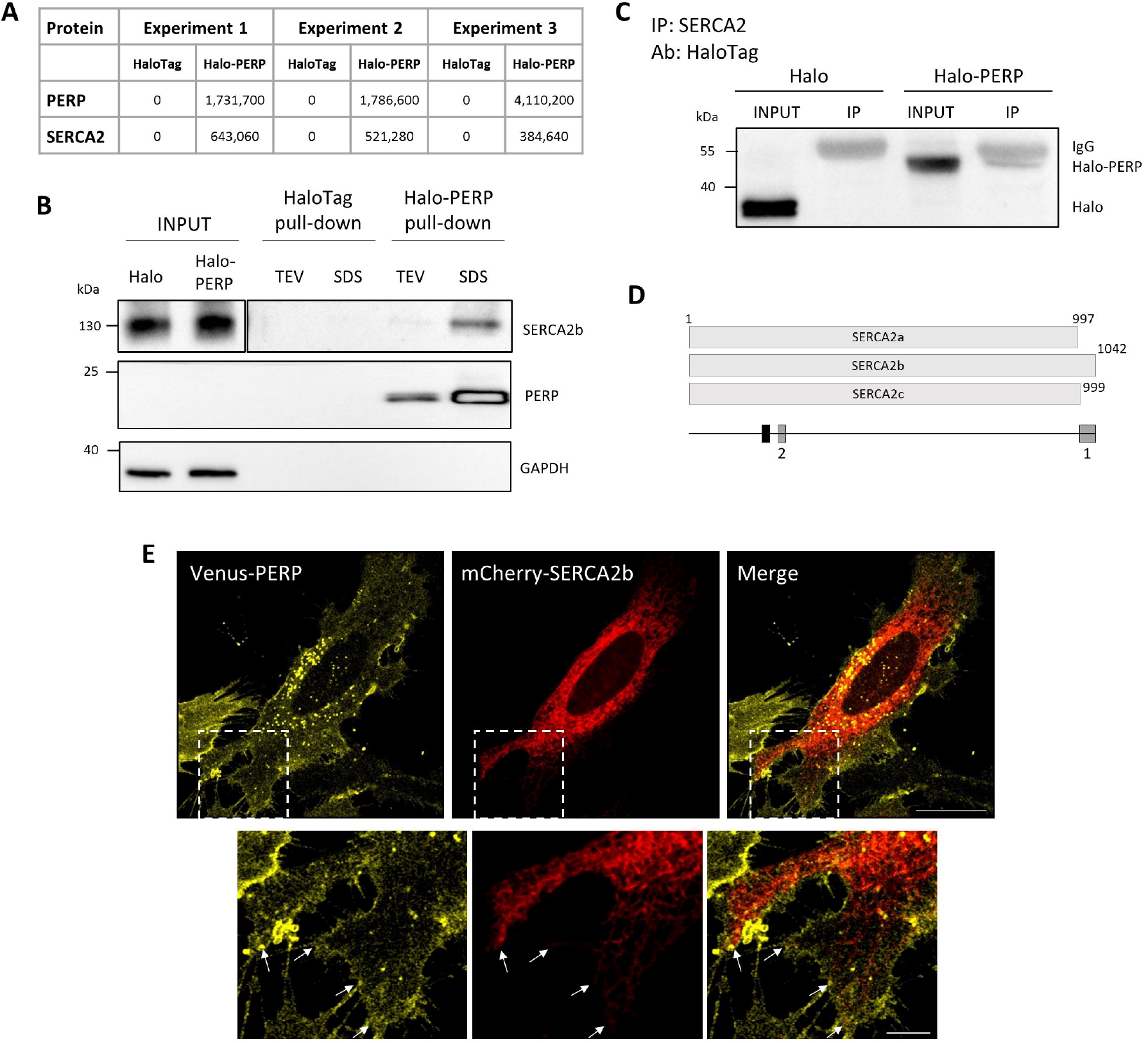
PERP interacts with SERCA2b at ER-PM junctions. **(A)** Protein interacting partners of Halo-PERP were isolated from Mel202 cells using the HaloTag Mammalian Pull-Down System and identified by mass spectrometry. Maxquant intensity values for SERCA2 and PERP in three independent experiments are shown. **(B)** The interaction of PERP and SERCA2b was confirmed in two independent HaloTag pull-down experiments by immunoblotting as described in the Methods (proteins eluted from the resin by TEV enzymatic cleavage followed by a successive SDS-based elution). Figure shows two different exposures of SERCA2b panel to allow visualisation of lower intensity bands). **(C)** PERP-SERCA2 complex formation was validated by SERCA2 IP in Mel202 cells expressing HaloTag or Halo-PERP using a HaloTag antibody. **(D)** Diagram of SERCA2a-c proteins, indicating the relative positions of the peptide identified by mass spectrometry (black) and the antibody immunogen sites (grey) used for validation of the SERCA2b-PERP interaction. Antibody 1 was used in HaloTag pull down validation **(B)** and antibody 2 was used for IP of SERCA2 **(C). (E)** Super resolution images of HeLa BAC Venus-PERP cells co-expressing mCherry-SERCA2b, junctions between the ER and PM shown by arrows. Scale bar 20 μm in full image and 5 μm in zoom panel.

To further validate the interaction between PERP and SERCA2, HaloTag and Halo-PERP were expressed in Mel202 cells and SERCA2 interacting proteins were isolated by immunoprecipitation (IP). The specific isolation of Halo-PERP was confirmed by Western blot using a HaloTag antibody (Fig. 1 C). Together these data identified and validated a consistent interaction between PERP and the ER Ca^2+^ pump SERCA2b (Fig. 1 D).

We next aimed to characterise the localization of the PERP-SERCA2b interaction. To this end, mCherry-SERCA2b was co-expressed in HeLa BAC Venus-PERP cells, which stably express Venus-PERP at physiological levels (Fig. 1 E). Venus-PERP localized to the PM and secretory pathway and mCherry-SERCA2b localized broadly across the ER including the cortical ER, which lies adjacent to the PM. Notably, PERP was present in regions of the PM in close co-localization with the SERCA2b-containing ER (shown by arrows).

The findings highlighted a novel interaction between the apoptosis effector protein PERP and the ER Ca^2+^ pump SERCA2b, which likely occurs at ER-PM points of contact.

### PERP protein is upregulated during ER stress independent of p53 transcriptional regulation

Since SERCA2b is known to be induced in response to ER stress^14^ and we have revealed an interaction between PERP and SERCA2b, we next explored the possibility that PERP may also respond to ER stress. Brefeldin A (BFA) prevents the export of proteins from the Golgi apparatus, leading to the accumulation of proteins in the ER and activating ER stress response pathways. Prolonged exposure to BFA induces caspase-dependent apoptosis.^30^ During conditions of ER stress, cells selectively transcribe genes with regulatory ER stress response elements in their promoters, including SERCA2b.^14^ BFA significantly increased the relative levels of SERCA2b mRNA from 2 h post-treatment (Fig. 2 A), followed by an increase in SERCA2b protein levels from 8 h post-treatment (Fig. 2 B).

**Figure 2.**
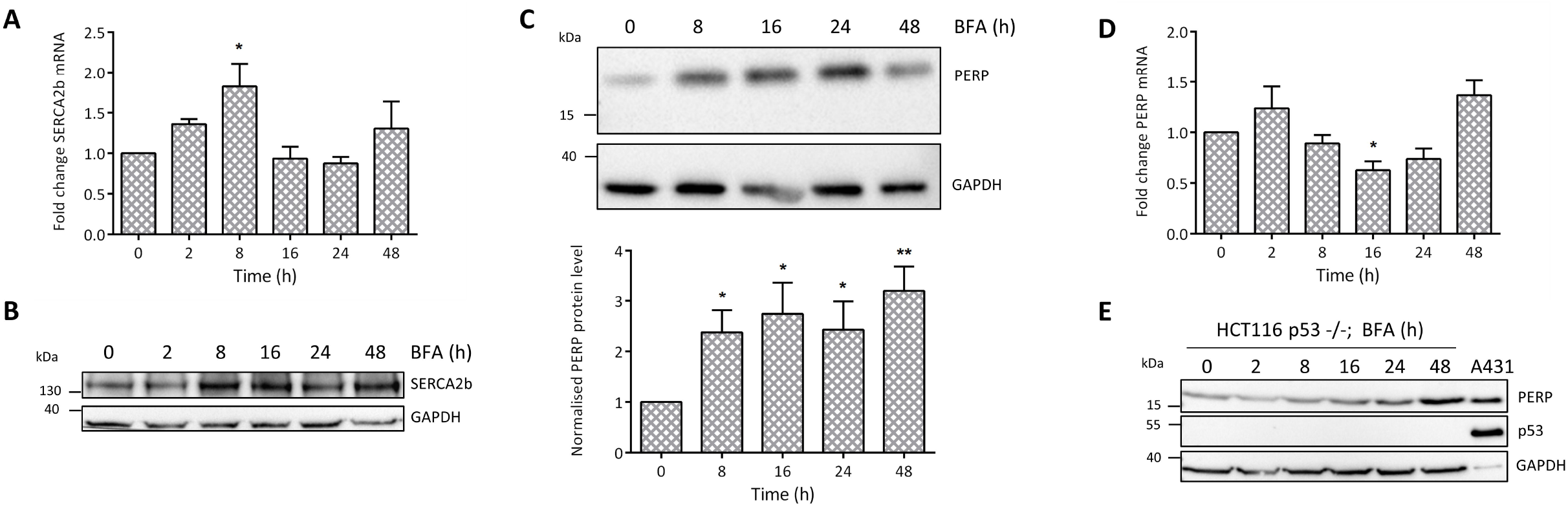
PERP protein is stabilized by ER stress, independent of p53 transcriptional regulation. HCT116 cells were treated with 1 μg/ml BFA for the indicated time points and changes to the levels of **(A)** SERCA2b mRNA (n=4, one-way ANOVA, F=3.442, p=0.0239*), **(B)** SERCA2b protein (n=7), **(C)** PERP protein (n=7, one-way ANOVA, F= 3.025, p= 0.0329*) and **(D)** PERP mRNA (n=4, one-way ANOVA, F=5.297, p=0.0036**) were detected by RT-PCR or Western blot and normalised to the level of GAPDH. **(E)** HCT116 p53−/− cells were treated with 1 μg/ml BFA and the response of PERP protein levels was detected by immunoblotting.

HCT116 cells were used to study the response of endogenous PERP to BFA-induced ER stress and treatment with BFA for 8, 16, 24 and 48 h significantly increased the protein levels of PERP (Fig. 2 C). We next wanted to determine whether the increase in PERP during ER stress was due to an increase in transcription. HCT116 cells were treated with BFA and PERP mRNA levels were measured by RT-PCR over time. BFA-induced ER stress led to no significant increase in the mRNA levels of PERP, which actually significantly decreased at 16 h post-treatment (Fig. 2 D). These findings were also consistent with those obtained in Mel202 cells (data not shown).

HCT116 p53−/− cells have significant deletions in exons 1-3 of the p53 gene and produce p53 protein that is defective in its ability to bind to DNA.^31^ Similarly to wild type cells, HCT116 p53−/− cells upregulated PERP protein in response to prolonged ER stress (Fig. 2 E), with no significant changes to PERP mRNA levels (data not shown). This finding suggested that p53 activity is not required for PERP to respond during conditions of ER stress. Furthermore, analysis of the PERP promoter region found no currently characterised ER stress response element in the first 1000 bases before exon one. This supported our finding that PERP is not transcriptionally regulated during conditions of ER stress.

In summary, the data presented here showed that although SERCA2b expression increased by transcription during conditions of ER stress, PERP protein levels increased via post-translational regulation in a p53-independent manner.

### PERP accumulates at the PM in response to ER stress due to a reduction in its turnover

PERP localization in response to ER stress was next assessed by super resolution live cell microscopy. Venus-PERP localized non-homogenously across the PM of non-treated cells and its levels were highest in regions of the membrane that formed contacts with neighbouring cells (Fig. 3 A). Treatment with BFA for 16 h induced the accumulation of Venus-PERP across the cell surface and within ER tubules, with the formation of ER vacuoles due to the prolonged induction of ER stress. The accumulation of ER-localized Venus-PERP was likely due to the BFA-induced inhibition of Golgi-PM trafficking and the retrograde transport of Golgi proteins into the ER. This finding confirmed that PERP reaches the PM via the classical secretory pathway.

**Figure 3.**
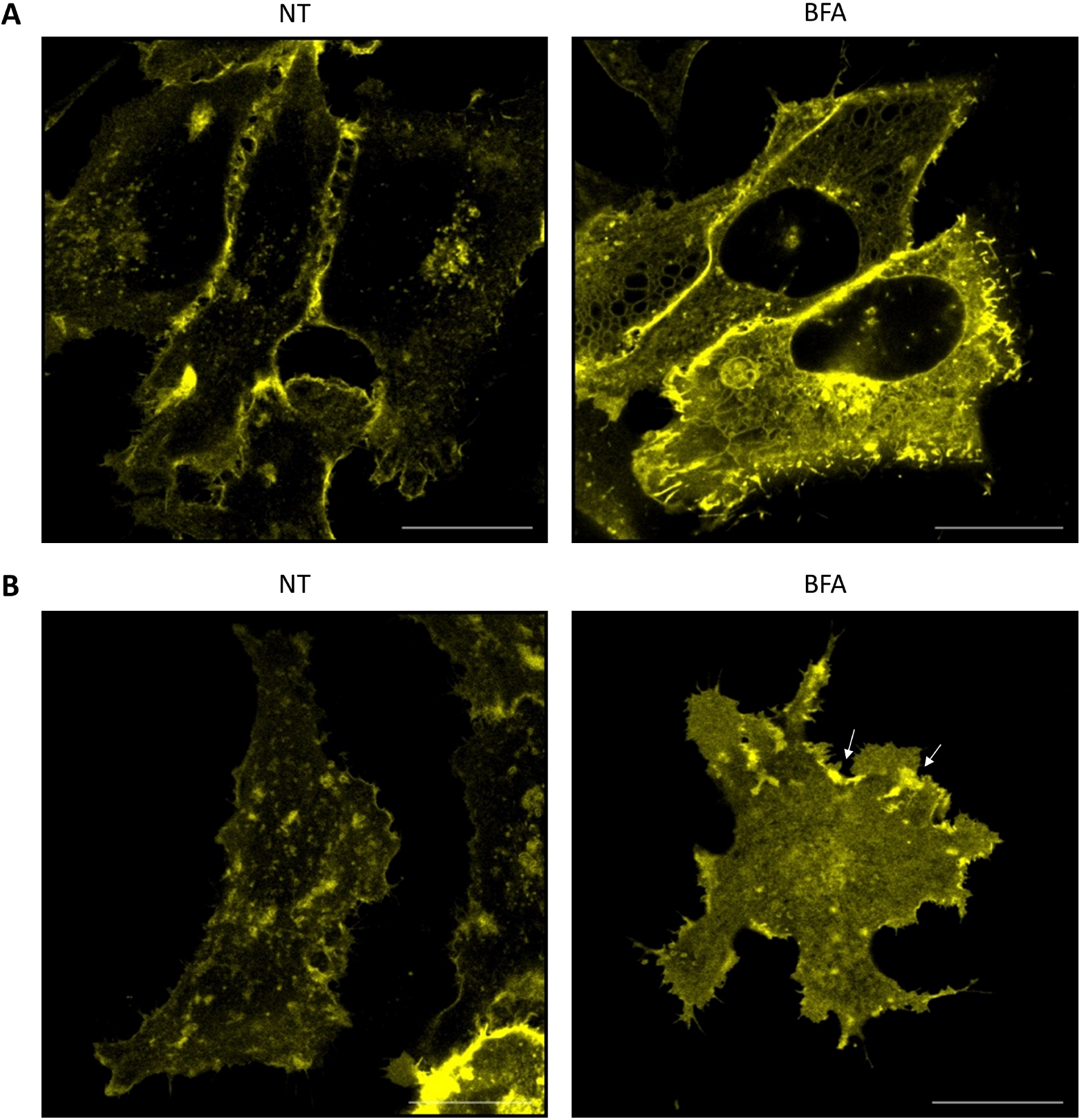
PERP accumulates at plasma membrane puncta during ER stress. **(A and B)** Super resolution images of HeLa BAC Venus-PERP cells at the **(A)** z-stack centre and **(B)** base of the cell, either non-treated (NT) or BFA-treated (16 h). Scale bars 20 μm.

The distribution of PERP at the PM during conditions of ER stress was analysed at the adherent base of the cell (Fig. 3 B). Venus-PERP localized to distinct PM islands in nontreated cells and significant areas of the membrane were PERP protein-free. In response to BFA treatment, Venus-PERP displayed a more homogeneous distribution across the PM and formed intense puncta at the cell periphery (shown by arrows). Significantly, since BFA inhibits Golgi-PM trafficking, the accumulation of Venus-PERP at the PM was not due to the delivery of newly synthesised protein. Therefore, the increase in the PM levels of PERP under ER stress conditions was likely due to a reduction in its translocation from the PM and in its subsequent degradation.

Together these findings indicated that PERP accumulates at the PM during ER stress via an increase in its protein stability and a reduction in its turnover.

### PERP protein is actively degraded by the lysosome

We therefore next characterised the stability and degradation of PERP protein. HCT116 cells were treated with the protein synthesis inhibitor cycloheximide (CHX) and the protein levels of PERP were detected over time (Fig. 4 A). p53, used as a positive control for protein synthesis inhibition, decreased from 2 h post-treatment due to its short half-life. The protein levels of PERP remained constant at 2 h post-CHX treatment followed by a significant decrease from 4 h post-treatment. Remarkably, PERP protein levels were reduced by 50% after 4 h of protein synthesis inhibition.

**Figure 4.**
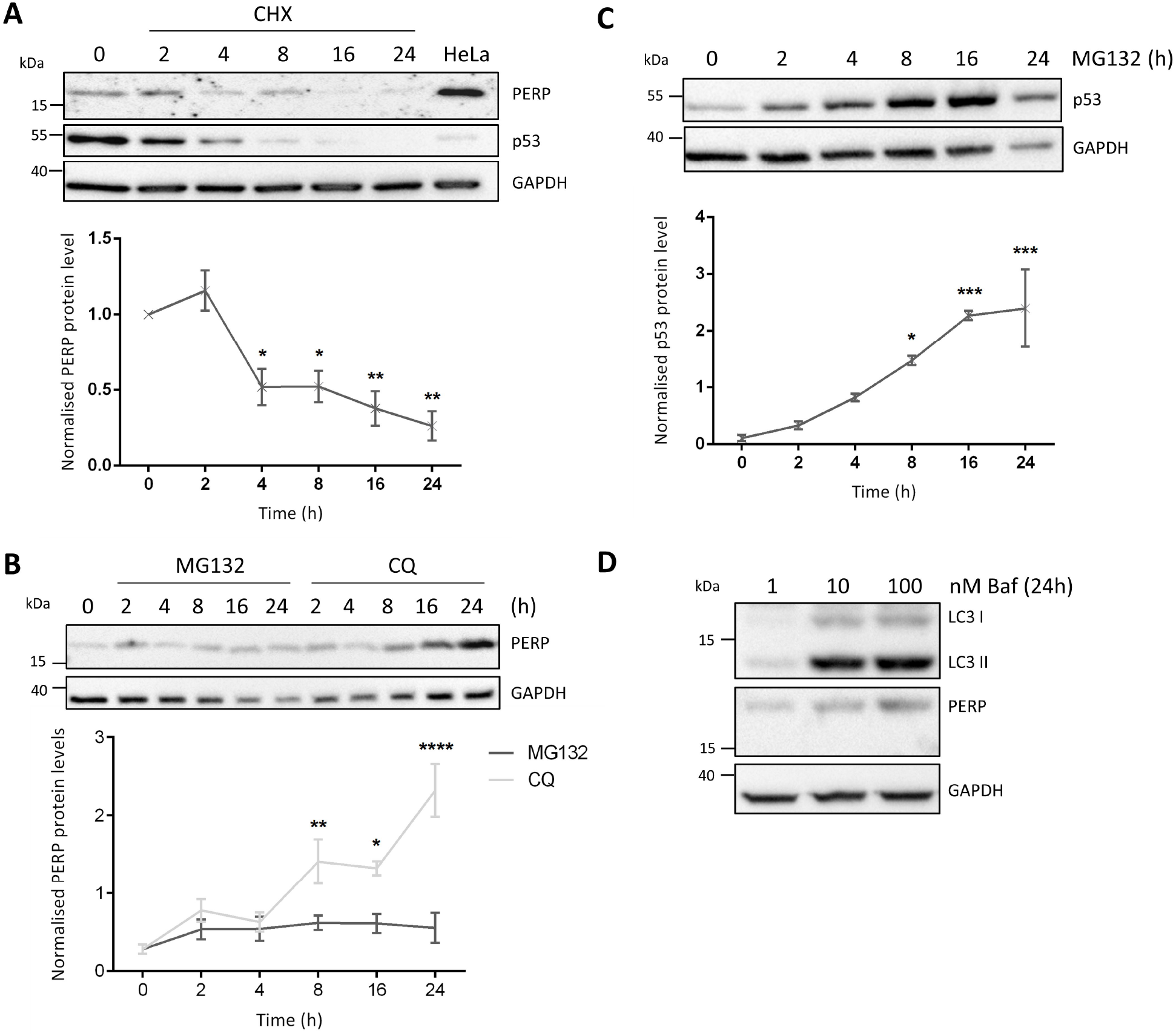
PERP protein is actively degraded by the lysosome. **(A)** HCT116 cells were treated with 30 μg/ml CHX for the indicated time points and the protein levels of PERP and p53 were detected by immunoblotting. Histogram shows PERP protein levels normalised to GAPDH. One-way ANOVA, n=3, F=12.63, p=0.0003***. **(B)** HCT116 cells were treated with 20 μM MG132 or 100 μM CQ for the indicated time points and PERP protein levels were detected by Western blot. Graph represents PERP protein levels normalised to GAPDH. One-way ANOVA, n=3, MG132 PERP: F=0.8849, p=0.5204 ns; CQ PERP: F=13.03, p=0.0002***. **(C)** HCT116 cells were treated with 20 μM MG132 and the protein levels of p53 were detected by immunoblotting and normalised to the level of GAPDH. One-way ANOVA, n=3, F=11.58, p=0.0003***. **(D)** HCT116 cells were treated with 1, 10 and 100 nM of bafilomycin (Baf) and the protein levels of LC3B and PERP were detected 24 h post-treatment by immunoblotting.

We also characterised the pathway that mediates the degradation of PERP protein. HCT116 cells were treated with the proteasome inhibitor MG132 or the lysosome inhibitor chloroquine (CQ) and cell lysates were collected for up to 24 h (Fig. 4 B). Inhibition of the proteasome degradation pathway led to no significant changes to PERP protein levels. However, PERP protein levels progressively increased from 2 h of lysosome inhibition, with a significant increase from 8 h post-treatment. This suggested that PERP protein undergoes lysosomal degradation. The protein levels of p53 were used as a positive control for proteasome inhibition by MG132; p53 increased from 2 h post-MG132 treatment with a significant increase from 8 h post-treatment (Fig. 4 C). PERP lysosomal degradation was confirmed using a second lysosome inhibitor bafilomycin (Baf) and the protein levels of both PERP and the autophagy marker LC3II increased in a dose-dependent manner 24 h after treatment (Fig. 4 D).

Together this data showed that PERP protein is actively turned-over in healthy cells by a pathway requiring lysosomal function.

### Upregulation of PERP by sustained autophagy precedes apoptosis induction

Multiple ER stress signalling pathways regulate autophagy and so we next characterised the response of autophagy markers LC3B and p62 to BFA (Fig. 5 A and B). BFA consistently increased the levels of LC3II from 2 h of ER stress induction and at 16, 24 and 48 h posttreatment LC3II increased beyond the dynamic range required for quantification compared to the non-treated sample. This confirmed that the number of autophagosomes progressively increased over time in response to BFA-induced ER stress.

**Figure 5.**
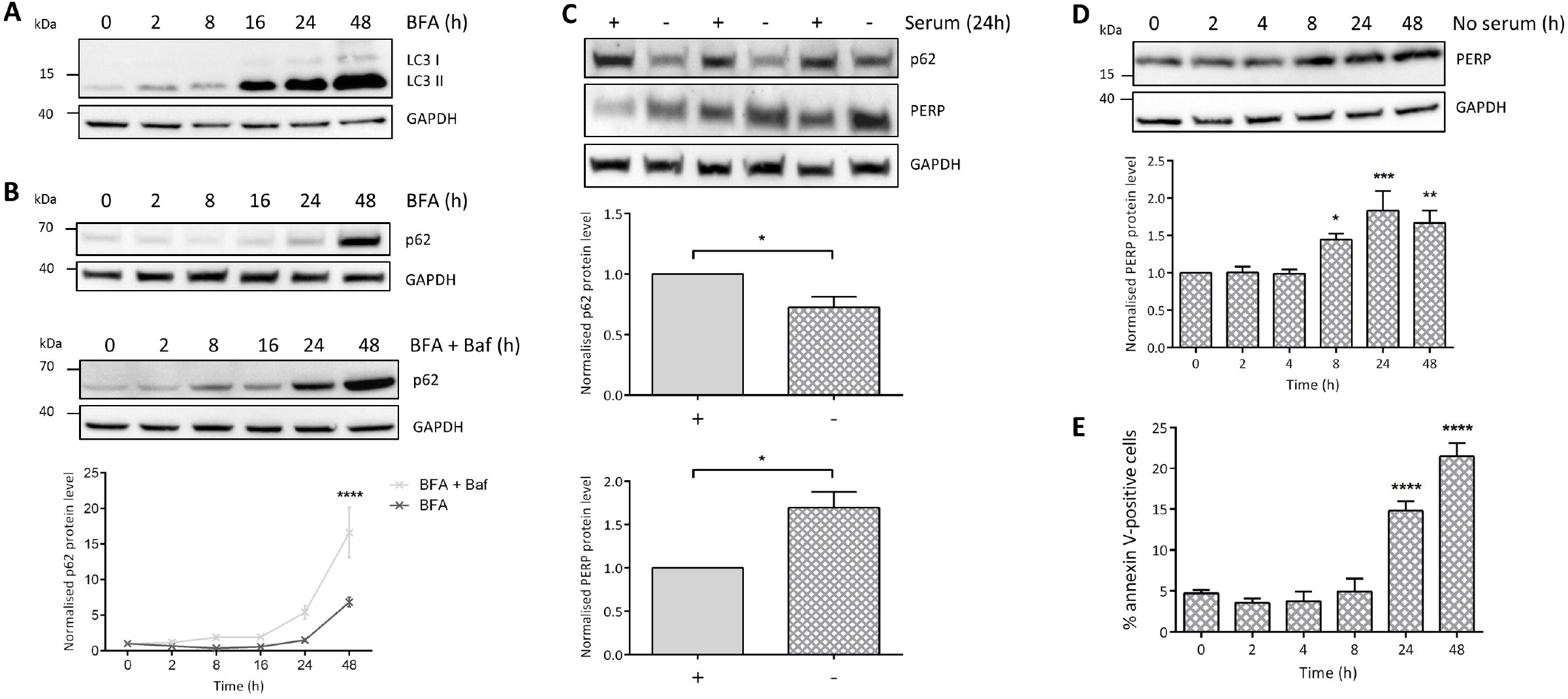
PERP is selectively upregulated in response to sustained autophagy induction, correlating with apoptosis. **(A)** Protein levels of autophagy marker LC3B in HCT116 cells treated with 1 μg/ml BFA. **(B)** Autophagy flux analysis of p62 levels in HCT116 cells treated with 1 μg/ml BFA in the presence and absence of 10 mM Baf. Protein levels quantified by densitometry and normalised to the level of GAPDH. Two-way ANOVA with Bonferroni post hoc test, n=3. **(C)** HCT116 cells were cultured in the presence (+) or absence (-) of serum for 24 h and p62 and PERP protein levels were detected and normalised to the level of GAPDH. Blot from one independent experiment shown, performed in technical triplicate. Student’s t-test, n=3, p62: p=0.0361*, PERP: p=0.02*. **(D)** HCT116 cells were serum starved for the indicated time points and PERP protein levels were detected by Western blot and normalised to the level of GAPDH. One-way ANOVA, n=5, F=7.426, p=0.0003***. **(E)** HCT116 cells were serum-starved for the indicated time points and apoptosis induction was measured by flow cytometry using an Alexa Fluor 647 annexin V conjugate. One-way ANOVA, n=4, F=41.00, p<0.0001****.

The protein levels of the autophagy marker p62 were used for autophagy flux analysis. BFA decreased the levels of p62 at 2 and 8 h, followed by an increase until a peak level at 48 h post-treatment. To determine whether the changes to LC3II and p62 protein levels in response to BFA were due to an increase in autophagy induction or an impairment in lysosomal degradation, HCT116 cells were pre-treated with Baf followed by BFA for the indicated time points (Fig. 5 B). Normalised p62 levels increased beyond that of the BFA only treated cells at all time points tested, which suggested that the increase in LC3II and p62 in response to BFA was due to an increase in autophagy induction rather than lysosomal inhibition.

We next determined the effect of autophagy induction on the protein levels of PERP. Firstly, HCT116 cells were serum starved for 24 h and the protein levels of autophagy marker p62 and PERP were determined (Fig. 5 C); p62 was significantly decreased confirming autophagy induction as characterised in this cell type elsewhere.^32^ In addition, PERP protein levels were significantly increased suggesting that PERP responds to autophagy induction.

We therefore next characterised the response of PERP to autophagy induction over time. HCT116 cells were serum starved for 2-48 h and the response of PERP was detected by immunoblotting. PERP protein levels remained constant for up to 4 h of serum starvation but significantly increased at 8, 24 and 48 h post-autophagy induction (Fig. 5 D). The sustained activation of autophagy promotes cell death and so the impact of serum starvation on apoptosis was determined over time by annexin V staining and flow cytometry; apoptosis was significantly induced from 24 h of serum starvation (Fig. 5 E).

Together our findings showed that during prolonged autophagy induction, such as that mediated by ER stress or serum starvation, PERP protein is upregulated before the switch to apoptosis induction.

### Increased SERCA2b leads to stabilization of PERP protein and induces apoptosis

Increased levels of SERCA2 are known to induce caspase-dependent apoptosis due to ER Ca^2+^ overload.^25,26^ To determine whether PERP is involved in SERCA2b-mediated apoptosis, mCherry-SERCA2b was expressed in Mel202 cells and changes to PERP protein and mRNA levels were detected (Fig. 6 A and B). At 24 h post-transfection, mCherry-SERCA2b was expressed at low levels and no changes to the mRNA or protein levels of PERP were observed. However, at 48 and 72 h post-transfection, both endogenous SERCA2b and mCherry-SERCA2b were increased compared to the control cells and this correlated with a significant increase in the protein levels of PERP. Despite this, expression of mCherry-SERCA2b for up to 72 h had no effect on the mRNA levels of PERP (Fig. 6 B). This suggested that increased expression of SERCA2b stabilizes PERP protein.

**Figure 6.**
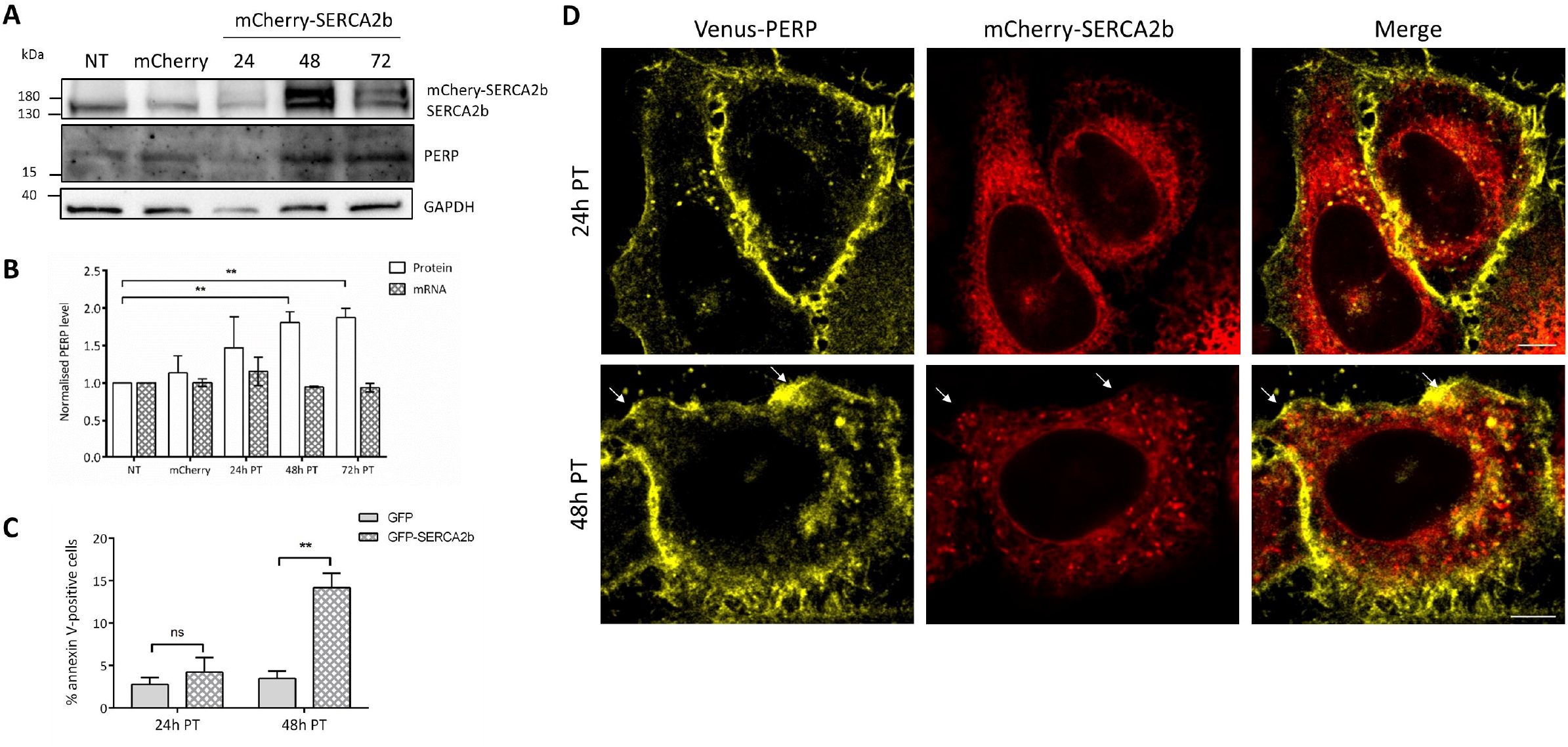
SERCA2b expression correlates with PERP protein stabilization and apoptosis induction. **(A)** mCherry (24 h) or mCherry-SERCA2b were expressed in Mel202 cells and PERP protein levels were detected by Western blot. **(B)** Graph shows PERP mRNA and protein levels normalised to the level of GAPDH in Mel202 cells expressing mCherry and mCherry-SERCA2b. Student’s t-test, n=3, 48 h PT p=0.005**; 72 h PT p=0.0017**. **(C)** The percentage of apoptotic Mel202 cells expressing eGFP and eGFP-SERCA2b was measured at 24 and 48 h post-transfection determined by flow cytometry using Alexa Fluor 647 annexin V. Student’s t-test, n=3, 24 h PT: p=0.4924 ns, 48 h PT: p=0.0048**. **(D)** Super resolution images of HeLa BAC Venus-PERP cells expressing mCherry-SERCA2b for 24 and 48 h, scale bars 20 μm.

The SERCA2b-mediated apoptosis in Mel202 cells was determined at both 24 and 48 h post-transfection by flow cytometry using annexin V (Fig. 6 C). At 24 h post-transfection with eGFP-SERCA2b there was no significant difference in the level of apoptosis compared to the eGFP expressing control cells. However, apoptosis was significantly induced at 48 h posttransfection with eGFP-SERCA2b. This showed that increased SERCA2b expression beyond the homeostatic threshold level at 48 h post-transfection mediated apoptosis. Furthermore, PERP was stabilized specifically during SERCA2b-mediated apoptosis.

Next, mCherry-SERCA2b was co-expressed in HeLa BAC Venus-PERP cells and images were taken at 24 and 48 h post-transfection (Fig. 6 D). At 24 h post-transfection the mCherry-SERCA2b-containing ER formed healthy tubules throughout the cell and subplasmalemmal region. However, at 48 h post-transfection mCherry-SERCA2b accumulated in intense puncta throughout the ER and cortical ER. Most notably, Venus-PERP was present in the regions of the PM that co-localized with the mCherry-SERCA2b puncta (shown by arrows in Fig 6 D).

Together, this data indicated that PERP protein is stabilized in the presence of apoptosis-inducing levels of SERCA2b expression.

### PERP and SERCA2b accumulate at ER-PM junctions in response to ER stress

Next, the effect of sustained ER stress on the interaction of PERP and SERCA2b was determined. HeLa BAC Venus-PERP cells co-expressing mCherry-SERCA2b were treated with BFA and images were taken 16 h post-treatment (Fig. 7 A). Imaging in the centre of the optical Z-stack of the cell showed co-distribution of Venus-PERP and mCherry-SERCA2b in the ER and vacuoles formed across the ER network. In addition, high levels of PERP across the cell surface were observed. Since the accumulation of PERP in the ER was potentially due to an indirect effect of the mechanism of BFA, the co-localization of Venus-PERP and mCherry-SERCA2b was determined at the base of the cell. Interestingly, mCherry-SERCA2b accumulated in intense puncta towards the PM in regions where Venus-PERP was also enriched (Fig. 7 B).

**Figure 7.**
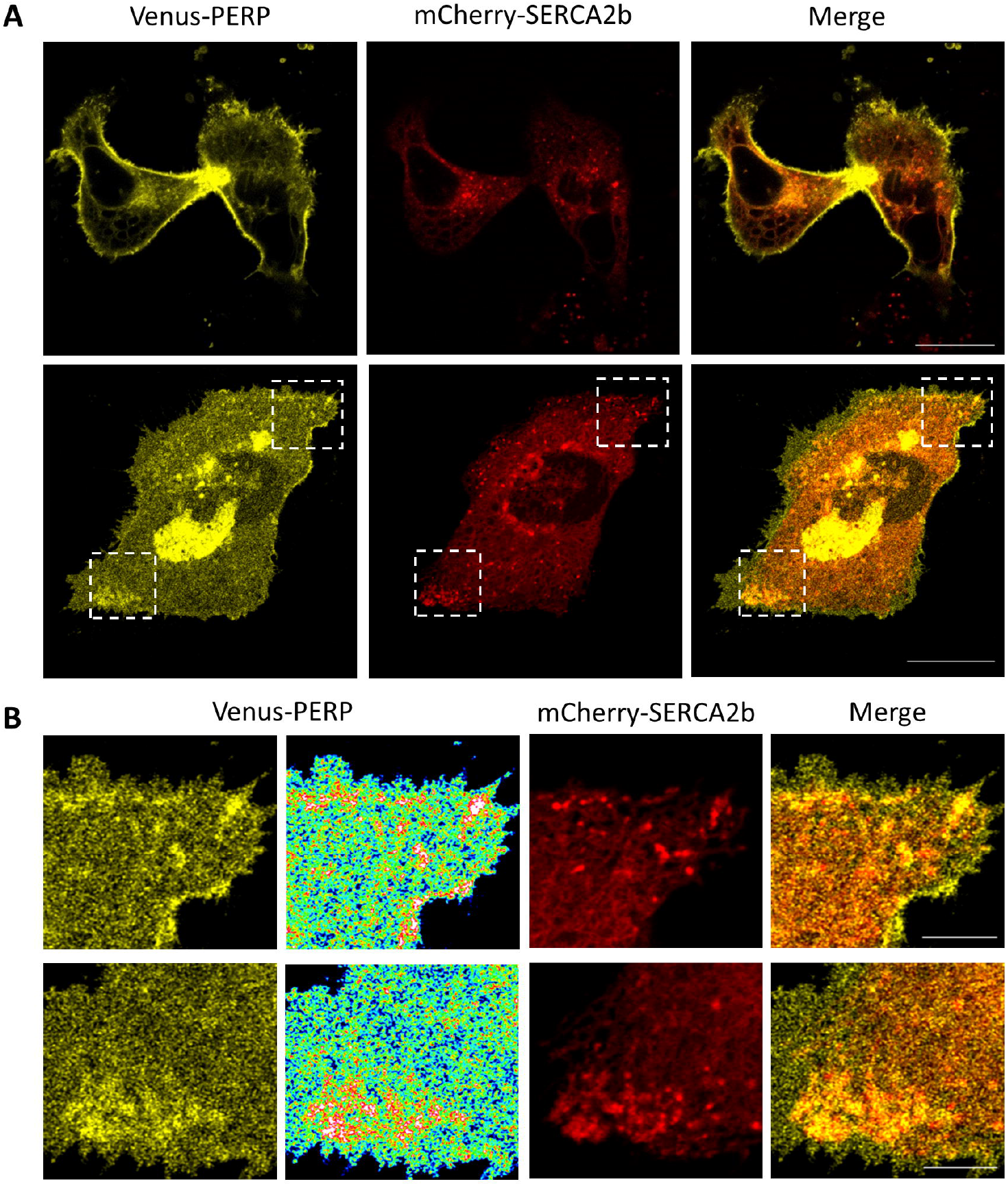
PERP increasingly interacts with SERCA2b during ER stress. **(A)** HeLa BAC Venus-PERP cells co-expressing mCherry-SERCA2b were treated with 1 μg/ml BFA for 16 h and colocalization was assessed at the mid and bottom optical section. Scale bars 20 μm. **(B)** Enlargement of boxed areas with rainbow pseudo colour-coded distribution of Venus-PERP. Scale bars 5 μm.

Taken together the findings indicate that during conditions of sustained ER stress, PERP accumulates at the PM due to a reduction in its lysosomal uptake and degradation, where it increasingly co-localizes with SERCA2b across ER-PM points of contact to mediate apoptosis.

## Discussion

The protein levels of the p53-regulated PM protein PERP are maintained at a threshold level in healthy cells.^6^ In this study, we showed that the threshold level of PERP protein is regulated through the action of a selective autophagy-lysosomal pathway. Therefore, PERP expression is precisely controlled by competing mechanisms that are sensitive to cellular state including, but not limited to, p53/p63-mediated transcription and autophagy-lysosomal protein degradation.

Using a cell-based model expressing physiological levels of PERP, we showed that PERP has a heterogeneous distribution across the PM of healthy cells with highest expression in regions of the PM that are engaged in cell-cell contacts. These areas are likely to be important for the cell adhesion function of PERP.^3,33^ In addition, the identification of an interaction between PERP and the ER-PM junctional protein SERCA2b highlighted, for the first time, localization of PERP to regions of the PM that come in close contact with the ER.

Expression of PERP above a threshold level influences cell fate towards the death outcome.^1,5–7^ Here, we showed that sustained autophagy, induced by both starvation and ER stress, increased PERP protein at the PM beyond the physiological threshold level and this correlated with apoptosis induction. Autophagy is also induced following *Salmonella* infection to aid pathogen clearance.^34,35^ Recently it was shown that PERP accumulates at the apical PM in response to *Salmonella* infection due to alterations in its uptake and degradation.^36^ We therefore propose that the upregulation of autophagy following infection promotes the stabilization of PERP at the PM.

Complex interactions between autophagy and apoptosis following cellular disturbances, such as ER stress, enable cells to dynamically regulate cell fate in a highly controlled manner.^37^ Our findings suggest that PERP is involved in the autophagy/apoptosis crosstalk; PERP is selectively upregulated at the PM following high levels of autophagy (starvation, ER stress, inflammation) where it directly engages its apoptotic machinery. In this scenario, PERP is protective against chronic autophagy.

Signals transduced across membrane contact sites via the formation of protein complexes and the transfer of molecules, such as Ca^2+^, regulate many cellular processes.^38^ SERCA2b is recruited to ER-PM junctions involved in SOCE where it is key to establishing Ca^2+^ homeostasis after oscillation.^11^ PERP lacks a conserved death domain and its precise mode of apoptosis induction from the PM is not understood.^7^ Here, we found that PERP is post-transcriptionally upregulated during SERCA2b-mediated apoptosis, conceivably through ER stress induced by dysregulation of luminal Ca^2+^ homeostasis.^12,25^ In addition, PERP and SERCA2b increasingly co-localize during chronic ER stress. PERP is the first identified PM-localized interactor of SERCA2b and we therefore propose that this interaction promotes the stabilization of SERCA2b in the cortical ER for sustained Ca^2+^ signalling events.

SERCA modulates the sensitivity to apoptosis and its Ca^2+^ pumping activity is regulated by competing pro-apoptotic and anti-apoptotic pathways.^39–42^ Apoptosis modulators, such as p53, activate SERCA2 to promote Ca^2+^-dependent apoptosis.^40^ Similarly, the PERP-SERCA2b interaction may mediate apoptosis by mitochondrial Ca^2+^ overload. This is supported by a study which showed that PERP induces apoptosis via an increase in mitochondrial membrane permeability and the release of cytochrome C in renal cells exposed to hypoxic injury.^43^

Our current findings provide the first mechanistic evidence of SERCA2 regulation and apoptosis induction at ER-PM junctions. The interaction of PERP and SERCA2b at junctions involved in SOCE may promote the sustained delivery of toxic levels of Ca^2+^ to the ER. However, PERP has a high sequence similarity with established Ca^2+^ channels and so it remains possible that PERP has Ca^2+^ conducting activity across the PM.^1^ The interaction between PERP and SERCA2b would therefore directly deliver extracellular Ca^2+^ into the ER for apoptosis.

This study has identified a novel crosstalk between the ER stress, autophagy and apoptosis pathways and has highlighted, for the first time, a mechanism of apoptosis regulation at ER-PM junctions. PERP-mediated destabilization of ER Ca^2+^ metabolism is likely to further induce both ER stress and autophagy responses and therefore amplify the stress signal to sway cell fate towards apoptosis.

## Acknowledgements

This work was supported by funding from the Humane Research Trust, UK and we are grateful to the laboratory of Javier García-Sancho for providing the SERCA2b constructs.

## Conflict of interest

All authors declare no conflicts of interest.

---

Supplementary Figure 1. **Halo-PERP plasma membrane localization and functional characterisation. (A)** Halo-PERP fusion protein localization was assessed in Mel202 cells using the fluorescent HaloTag TMRDirect ligand. Scale bar 20 μm. **(B)** Protein levels of endogenous PERP and p53 in Mel202 cells expressing HaloTag (24 h) or Halo-PERP for the indicated time points. One-way ANOVA, n=3, p53: F=13.39, p=0.0001***; PERP: F=17.12, p0.0001****. **(C)** The percentage of apoptotic Mel202 cells expressing HaloTag and Halo-PERP labelled with the R110 HaloTag fluorescent ligand 48 h post-transfection determined by flow cytometry using Alexa Fluor 647 annexin V. Student’s t-test, n=3, p=0.0026**.

