## Supplementary figures and images for "ER stress-linked autophagy stabilizes apoptosis effector PERP and triggers its co-localization with SERCA2b at ER-plasma membrane junctions"

### Supplementary Figure 1

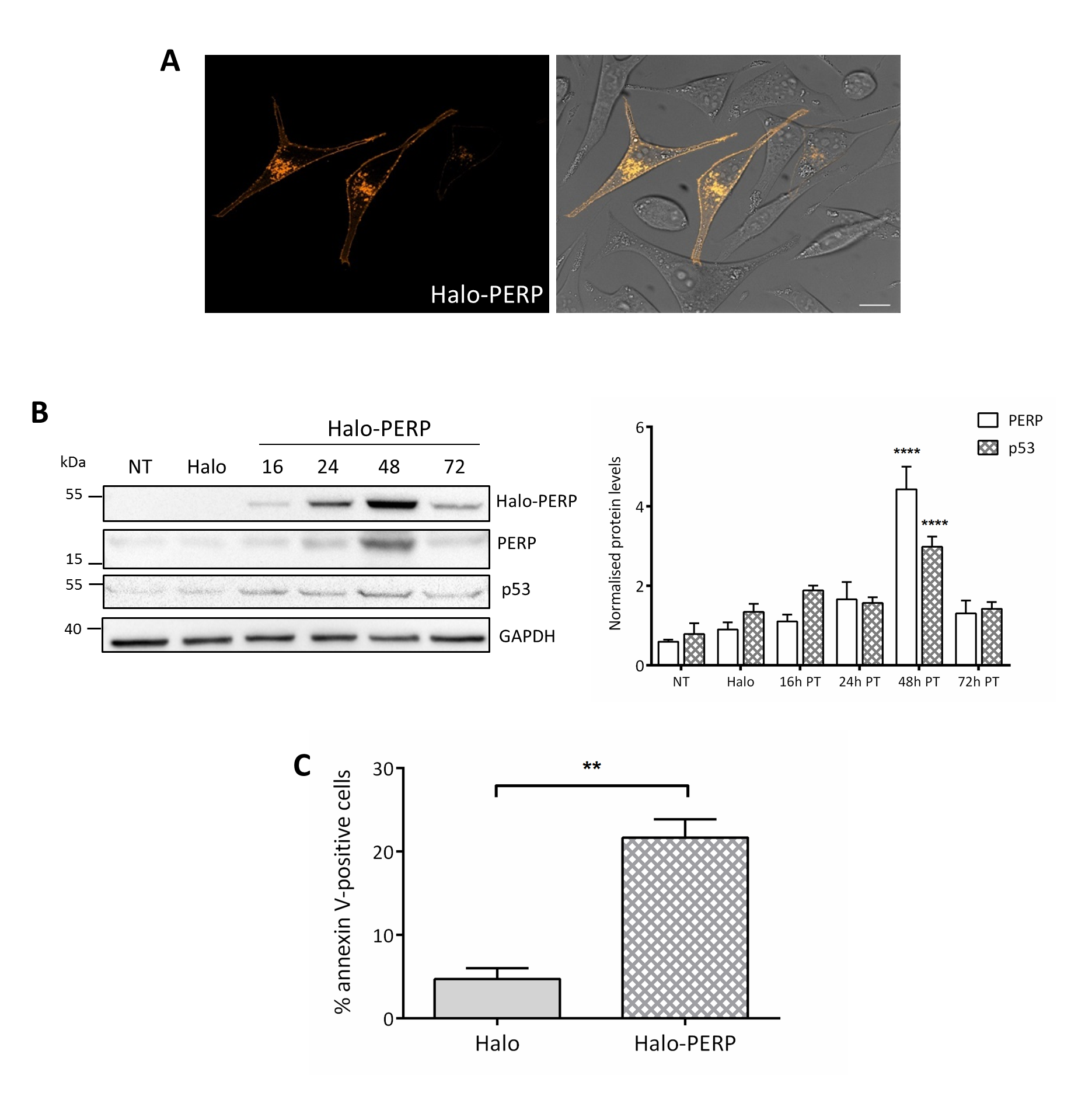
